# Sound disrupts sleep-associated brain oscillations in rodents according to its meaning

**DOI:** 10.1101/2021.09.29.462343

**Authors:** Philipp van Kronenberg, Linus Milinski, Zoë Kruschke, Livia de Hoz

## Abstract

Sleep is essential but poses a risk to the animal. Filtering acoustic information according to its relevance, a process generally known as sensory gating, is crucial during sleep to ensure a balance between rest and danger detection. The mechanisms of this sensory gating and its specificity are not understood. Here, we tested the effect that sounds of different meaning had on sleep-associated ongoing oscillations. We recorded EEG and EMG from mice during rapid-eye movement (REM) and non-REM (NREM) sleep while presenting sounds with or without behavioural relevance. We found that sound presentation per se, in the form of an unfamiliar neutral sound, elicited a weak or no change in the sleep-dependent EEG power during NREM and REM sleep. In contrast, the presentation of a sound previously conditioned in an aversive task, elicited a clear and fast decrease in the sleep-dependent EEG power during both sleep phases, suggesting a transition to lighter sleep without awakening. The observed changes generally weakened over training days and were not present in animals that failed to learn. Interestingly, the effect could be generalized to unfamiliar neutral sounds if presented following conditioned training, an effect that depended on sleep phase and sound type. The data demonstrate that sounds are differentially gated during sleep depending on their meaning and that this process is reflected in disruption of sleep-associated brain oscillations without an effect on behavioural arousal.

## Introduction

Sleep poses a risk to the animal due to the accompanying decreased behavioural response to sensory stimuli^1,2^ and the associated species-specific sleep posture^3^. This risk is an obvious necessary trade-off to carry essential physiological processes^4-7^ such as brain homeostasis/restoration processes^8-10^ or memory consolidation^11^. Sensory disconnection during sleep is not complete, however, and some continuation of sensory processing remains in order to detect behaviourally relevant stimuli. The mechanisms and specificity of this continued sensory processing are not understood. Here we explore the role of meaning in the effect that sound has on sleep-associated brain oscillations in the rodent brain.

Sensory processing of stimuli is reduced in the sleeping brain across modalities (olfactory^12,13^, visual^14^ and somatosensory^15,16^) with the exception of the processing of sounds by different auditory stations, including the primary cortex^17^. That sounds are treated differently from other stimuli is maybe not surprising since sounds, being detectable from far and around in a fast manner^18^, are valuable in the detection of approaching dangers. Here we tested the effect that neutral and danger-predicting sounds had on mouse sleep-associated ongoing brain oscillations during different sleep phases, in order to better understand the mechanisms and selectivity of sound processing during sleep. Other animals, especially cats have been used to investigate arousal thresholds^19^ and habituation^20,21^ during sleep, but we have little understanding about the role of sound meaning in these processes^22^.

We developed an auditory associative learning task, which we combined with sleep EEG recordings. During conditioning, a sound was associated with an aversive experience. This sound, as well control sounds of neutral meaning to the animal, were then presented to the mouse during sleep. Ongoing brain oscillations, measured through EEG, were the readout to determine if a sound had a visible effect on the sleeping brain. Sensory gating, the filtering of stimuli according to their relevance^23^, is crucial in our everyday life, and especially important to achieve a continuous sleep ^24^. Hence, sleep is an effective model to study sensory gating mechanisms. The combined behavioural/EEG paradigm is useful for the investigation of the circuit regulating sensory gating in the sleeping and, potentially, waking brain.

## Results

To study sound processing during sleep, we played meaningful and neutral sounds during different sleep phases to nine female C57BL/6J mice. First, during a 6-day long ‘exposure phase’, we recorded the EEG in sleeping mice for 1.5-2 hours per day. During this time, we played a fixed ‘pre-control’ pure tone at different times during sleep to assess the effect that a neutral sound has on different sleep phases. For the following 4-8 days, we then trained animals to escape an aversive stimulus (loud broadband noise plus air puff) that followed a conditioned frequency-modulated (FM) sound. Daily, during subsequent 1-2.5 hour-long sleep sessions, we recorded the EEG and EMG activity in freely moving mice to identify REM (rapid eye movement) and NREM (non REM) phases of sleep, and played the conditioned and an unfamiliar neutral FM sound (post-control sound), in increasing intensity (Fig. 5b), for 8 - seconds during either sleep phase. The post-control sound aimed at dissociating the effects of the conditioned sound that resulted from its valence as a predictor of punishment from those that resulted from its novelty.

### Validation of manual sleep phase classification

During the experiment, we used online visual inspection, based on conventional criteria in sleep scoring^25,26^, of the ongoing oscillations to categorize the current mental state as awake, REM, NREM, or unidentifiable. To ensure that this decision matched a quantifiable measure, we first analysed the recorded EEG. We found that the experimenter’s scoring paralleled the spectrogram of the parietal EEG (Fig. 1a). Phases that had been categorized as NREM sleep were indeed characterized by synchronized slow wave activity (delta frequency band, 0-4 Hz) and high amplitude irregular activity typical of NREM (Fig. 1a, orange trace). In contrast, phases categorized as REM sleep were characterized by fast and desynchronized activity (theta band, 6-10 Hz) with lower amplitude and high regularity (Fig. 1a, purple trace) as described by Jouvet and collegues^27^. A quantification of both the theta/delta power ratios and the regularity of the amplitude in 4-second long EEG time windows, clearly separated our subjectively classified NREM (orange) and REM (purple) into two well separated clusters (Fig. 1b). NREM sections had low theta power and high delta power (Fig. 1c) and the peaks’ standard deviation was high, as expected from the typical NREM activity. For sections scored as REM sleep, we found high theta and low delta activity (Fig. 1c) together with a low variability in peak amplitude. To conclude, the quantification of the EEG signal validated the subjective characterization of NREM and REM sleep performed during the experiment.

**Figure 1.**
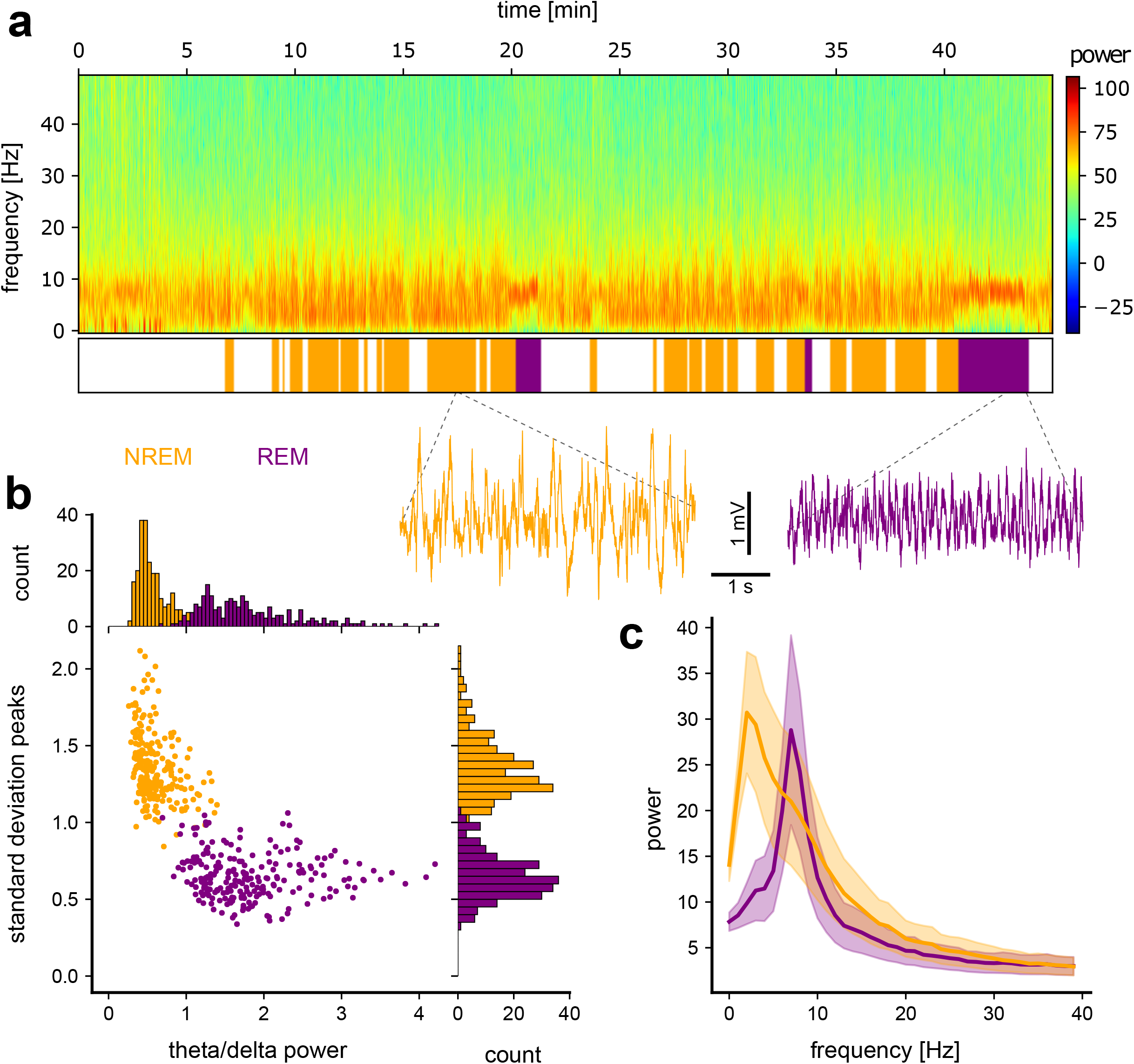
Sleep stage characterization. **(a)** Top: Power spectrogram of an example parietal EEG recording including sleep. Middle: Visual sleep scoring (based on EEG+EMG; awake = white, NREM = orange, REM = purple) of the spectrogram shown above. Bottom: two example EEG traces taken from a typical NREM (orange) / REM (purple) sleep bout. **(b)** The scatter plot depicts the power ratio in the theta/delta frequency bands against the peak-to-peak variability (standard deviation). Data from undisturbed sleep. The histograms indicate the distinct distributions for each measure respectively for NREM and REM sleep. (N (animals) = 11, n (sleep sections) = 246 (NREM), 228 (REM)) **(c)** Mean spectral power distribution for NREM and REM sleeps. (N (animals) = 11, n (sleep sections) = 246 (NREM), 228 (REM))

### The conditioned sound elicits reductions in state-dependent EEG power during NREM and REM sleep

To quantify changes in the EEG signal elicited by sound presentation we analysed the power in the delta frequency band (1-4 Hz) for NREM sleep and in the theta frequency band (6-10 Hz) for REM sleep. The power was analysed for each individual EEG trace over 24 seconds (eight seconds before sound onset, eight seconds during sound presentation at increasing intensity, and eight seconds after the sound ceased). Visual inspection made it evident that there were differences in the effect that pre-control, conditioned and post-control sound had on both NREM and REM sleep (Fig. 2a, 2c). In a raw EEG example trace recorded during NREM, one can observe that the typical 1-4 Hz delta activity, which had not been influenced by the pre-control sound (Fig. 2a top), was disrupted by both the post-control and conditioned sounds (Fig. 2a mid and bottom). This difference in effects is also evident in the quantification of the mean spectral power over the first 4 days of conditioning, with superimposed mean for the last 4-days of exposure. The power in the delta range (0-4 Hz) did not change with pre-control sound presentation, but decreased drastically soon after the conditioned and post-control sounds were played (Fig. 2b). It only began to recover several seconds after the sound ceased. A 2-way ANOVA on sound type x time for the block of sound presentation during NREM, revealed a main effect of sound (F(2,120) = 40.24, p<0.001), time-block (F(7,120) = 15.11, p<0.001) and interaction between sound and time (F(14,120) = 2.16, p=0.0133). Only animals that learned the task were included here (over 80% avoidance on the second day and at least 15 conditioning trials, Fig. 2b, inset). The slight drop in power towards the end of the pre-control sound presentation coincided with the loudest intensities of our sound sequence. In contrast, the drop in power was stronger and already very prominent for post-control and conditioned sound during the lower intensity repetitions of the sound sequence, at intensities somewhere between 30 and 40 dB SPL. That the post-control but not the pre-control sound elicited a change comparable to that of the conditioned sound suggests some form of generalization to the conditioned sound in the sensitivity to sound during NREM.

**Figure 2.**
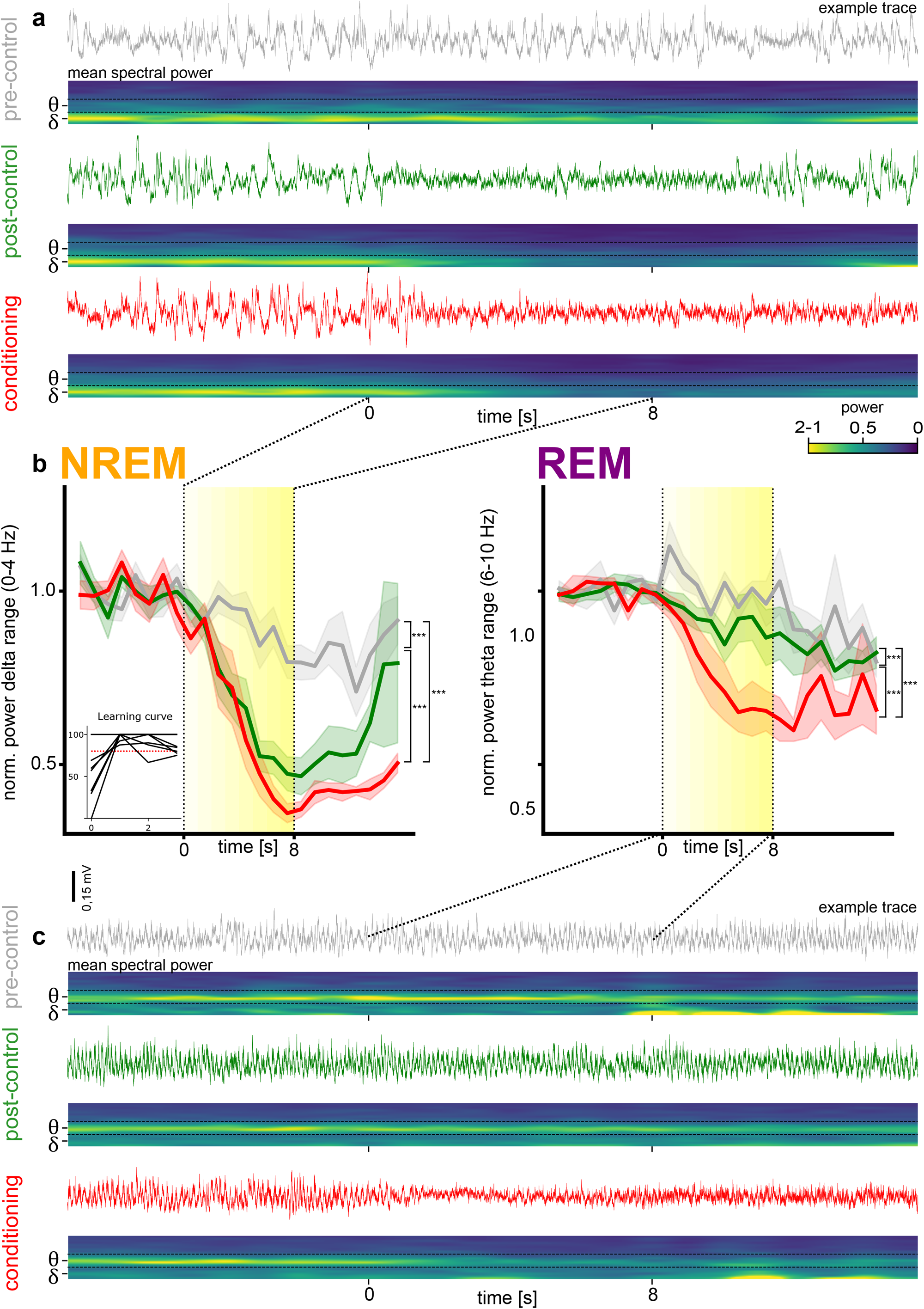
Sound-driven EEG power changes during NREM and REM sleep. **(a) NREM. Top traces:** Example EEG traces and mean power spectrograms for each experimental sound during NREM sleep. **Bottom spectrograms:** Mean power spectrograms representing the mean spectral power over all animals and sleep sections for each sound type played. δ = Delta frequency band (0-4 Hz), θ = Theta frequency band (6-10 Hz). Time points 0 and 8 mark the beginning and end of sound presentation respectively. **(b)** The mean normalized EEG power in the delta (for NREM) and theta (for REM) ranges over time for learner animals (> 80% correct trials on second day and > 15 trials in total) as the pre-control (grey), post-control (green), and conditioned (red) sounds were presented. Sound presentation is illustrated by a yellow block with sound intensity reflected in the colour intensity. In the figure, significance level is indicated as follows: * = p<0.05, ** = p<0.01, *** = p<0.001, (N = 6 (trials = 246 (NREM), 228 (REM)). **(c) REM**. Same as (A), but for REM sleep.

During REM, the pattern was somewhat different. As can be seen in the example traces (Fig. 2c), only the conditioned sound elicited a change in EEG pattern. This pattern persists in the spectral analysis of the mean 6-10 Hz theta activity during sound presentation over the first 4 days of conditioning. In the quantification graph of REM sleep for all three experimental sounds (Fig. 2b right), we confirmed the interesting disparity of the power changes to the post-control sound during NREM sleep. Only the conditioned sound elicited a strong power change during REM.

A 2-way ANOVA on sound type x time for the block of sound presentation during REM, revealed a main effect of sound (F(2,120) = 31.6, p<0.001) and time-block (F(7,120) = 3.46, p<0.0021) without interaction. Hence, animals showed higher generalization to the experimental sounds during NREM sleep compared to REM sleep.

Generally, power changes were weaker during REM sleep than during NREM sleep. The averaged changes during NREM sleep to the conditioned sound were dropping to a minimum of 36% of the original power, whereas for REM sleep 59% was the global minimum. The same was true for the other experimental sounds, which showed a fundamental difference between NREM and REM power changes. A 2-way ANOVA on sleep phase x sound type x time-block (first 8 seconds, 8 seconds during sound presentation and the 8 seconds afterwards), revealed a main effect of state (F(1,850) = 84.77, p<0.001), sound (F(2,850) = 82.92, p<0.001) and time-block (F(2,850) = 203.64, p<0.001) and interaction between state and sound (F(2,850) = 5.53, p=0.0041), state and time-block (F(2,850) = 21.64, p<0.001) and sound and time-block (F(4,850) = 21.36, p<0.001).

The sound presentation was occasionally accompanied by muscle twitches, which were visible in the EMG. After the sound ceased, however, the animals continued sleeping without awakening. Nevertheless, the change in EEG pattern to a more desynchronized pattern during NREM sleep and to a more irregular pattern during REM sleep, point in the direction of a more unstable/alert state of the brain.

Overall, the pre-control sound led to subtle reductions in power that were small and late relative to sound onset. In contrast, the conditioned sound elicited strong and fast reductions in state-dependent EEG power during both NREM and REM sleep. Interestingly, this effect was generalized to the post control sound, in a weaker version, only during NREM sleep.

### Power change effects strongest during early conditioning days

To assess whether the observed effects changed over the course of days as the sounds became more familiar, we analysed the sleep effects on a daily basis over the first 8 days of conditioning (Fig. 3a, b). We found a decreased effect for the conditioned sound during NREM sleep. A 2-way ANOVA on the day of sound presentation during NREM sleep during and after the conditioned sound was played (8 seconds during sound presentation and the 8 seconds afterwards), revealed a main effect of day (F(7,591) = 22.59, p<0.001). Yet, only days 7 and 8 were significantly different from all previous days, suggesting that the effect was initially robust. This pattern was not replicated for the post-control sound during NREM sleep. A 2-way ANOVA on the day of sound presentation during NREM sleep for the post-control sound revealed no effect of day F(7,559) = 2.02, p=0.0513. With no days being significantly different from each other.

**Figure 3.**
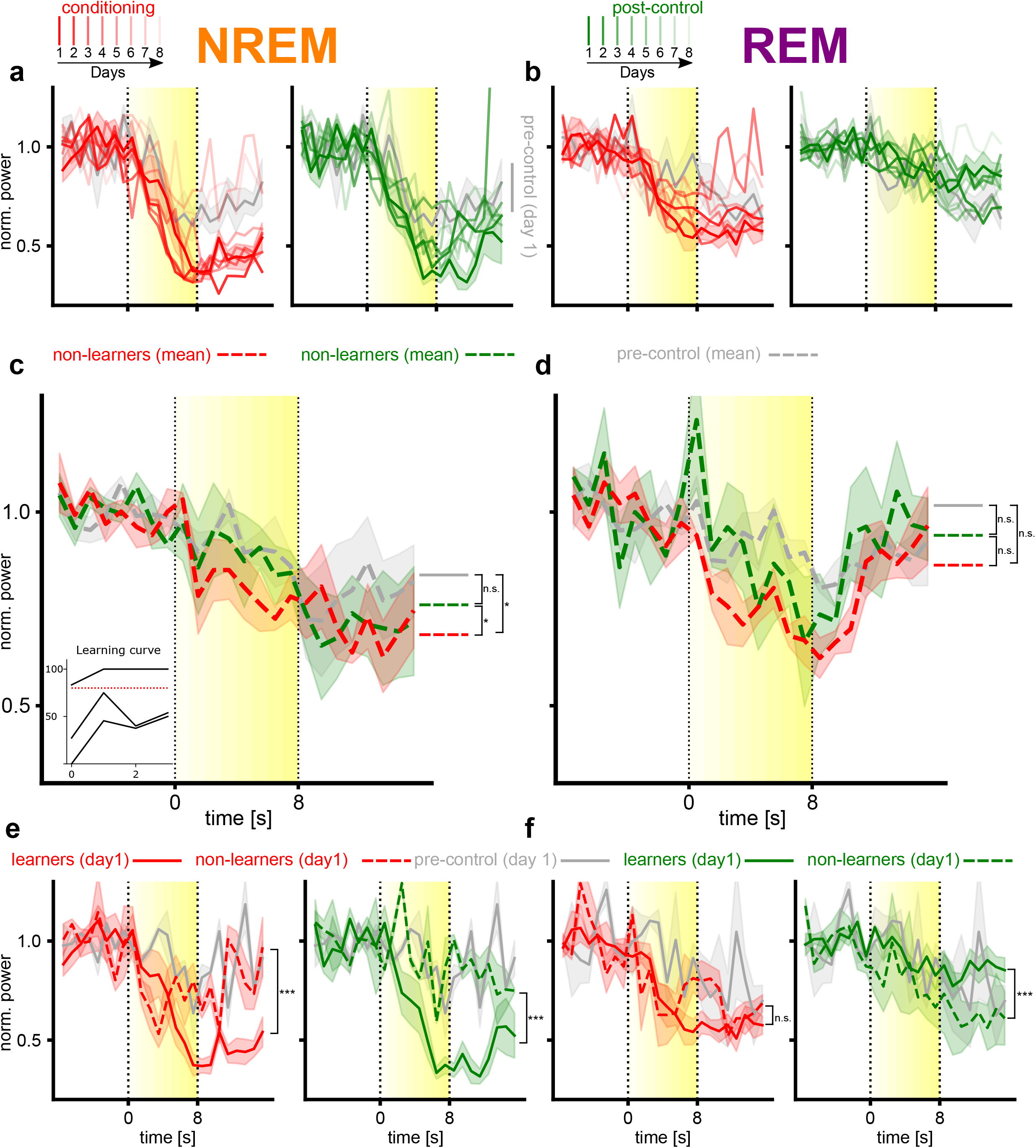
Effects of days and learning rate on EEG power changes. **(a-b)** Comparison of mean daily EEG power changes during **(a)** NREM sleep and **(b)** REM sleep over days for the conditioned (red) and post-control (green) sounds. Changes are measured over eight consecutive days with decreasing colour strength, starting with day 1, which is the first day of conditioning. Responses to pre-control sound on the first day serve as reference. (N=6) **(c-d)** Mean EEG power in **(c)** delta for NREM and **(d)** theta range for REM over time for non-learner animals (< 80% correct trials on second day or <15 trials in total) as the pre-control (grey), post-control (green), and conditioned (red) sounds were presented. Sound presentation is illustrated by a yellow block with sound intensity reflected in the colour intensity. In the figure, significance level is indicated as follows: * = p<0.05, ** = p<0.01, *** = p<0.001. (N = 3). **(e-f)** Comparison of EEG power changes during **(e)** NREM sleep and **(f)** REM sleep phases of the first recording session after conditioning. Learners (>= 80% correct trials, solid line) and non-learners (<80% correct trials, dashed line) are defined by second day behavioural performance in the conditioning paradigm. Responses to pre-control sound on the first day serve as reference.

During REM sleep, both sounds showed a main effect of day during an 2-way ANOVA with F(7,543) = 4.87, p<0.001 for the post-control and F(7,543) = 7.14, p<0.001 for the conditioned sound. The conditioned sound showed a decline in strength over days, with days 4 and 7 being significantly different from days 1-2 and day 4 being significantly different from days 5 and 8. The effect for the control sound, which anyway had a weak influence over REM sleep, is driven by the 8th day, which is significantly different from days 1, 2, 5 and 7.

Thus, the effect the conditioning sound had on ongoing EEG oscillations during sleep was weakened over days, a process that appeared earlier in REM than NREM.

### Non-learners show no distinct EEG changes to any experimental sound

Three out of the 11 mice trained did not learn to avoid the air-puff predicted by the conditioned sound or did not enter the corner sufficient times after the first experience with the conditioned sound. To study how non-learners differ from learners over the first four days of conditioning, we analysed the non-learners data exactly as we did for the learners in Fig 2b. Compared to learners, sound in non-learners elicited no clear effect in overall power (Fig. 3c, d). Nevertheless, EEG pattern changes during NREM and REM sleep for non-learners were significantly different from each other. A 2-way ANOVA on sleep state x sound type x time-block (first 8 seconds, 8 seconds during sound presentation and the 8 seconds afterwards), revealed a main effect of state (F(1,418) = 5.58, p=0.0186), sound (F(2,418) = 6.44, p=0.0018) and time (F(2,418) = 59.3, p<0.001) and an interaction between state and time-block (F(2,418) = 9.2, p<0.001). Additionally when looking at sound differences during NREM sleep in block two and three there was a main effect of sound (F(2,120) = 5.5, p=0.0052), driven by the conditioned sound which was significantly different from both pre-control (p=0.0122) and post-control (p=0.0146) sounds during multi-comparison. During the second and third block of REM sleep we did not find any effect of sound type (F(2,120) = 2.19, p=0.1159) or time in block (F(7,120) = 0.43, p=0.8848) in a 2-way ANOVA.

Hence, the responses of the non-learners were less specific and delayed compared to learners. The differences between power changes of learners and non-learners emphasize the importance of learning in the influence of sound meaning over sleep.

Interestingly, the behavioural performance on day two (i.e. whether animal was a ‘learner’ or ‘non-learner’) could be predicted based on the strength of the sound effects on day 1 on NREM (Fig. 3e) and for the post-control sound during REM (Fig. 3f)

Conducting a 2-way ANOVA we found a main effect of learning for the conditioned sound (F(1,142) = 16.91, p<0.001) and the control sound (F(1,126) = 31.27, p<0.001) during NREM sleep. During REM sleep we found no main effect for the conditioned sound (F(1,142) = 3.64,p=0.0583), only for the post-control sound (F(1,126) = 18.49, p<0.001). The learners showed strong EEG pattern changes during conditioned sound presentation during both NREM and REM, whereas for the non-learners weaker changes or different temporal curves could be seen. Learners also showed stronger power changes for the post-control sound during NREM and REM. Although, this power change was not comparable to the power changes to the conditioned sound during NREM and REM sleep from learning animals (Fig. 2b), because the change started comparatively late, only with the presentation of the loudest repetitions of the sound sequence.

### Sound generalization during NREM and REM sleep

To explore further the generalisation between conditioned and post-control FM sounds that we observed during NREM (Fig. 2b), we tested additional sounds with a spectrotemporally richer sound architecture. We trained two female C57BL/6J mice on the audio-terrace as before but unlike in the previous experiment, we used three, instead of one, post-control sounds. Additionally, in contrast to the experimental sounds previously used, we used sound clouds: a concatenation of pure tones pseudo randomly selected between set frequency borders (pre-control: 3000-4662 Hz, post-control 1: 4811-7478 Hz, post-control 2: 7718-11995 Hz, post-control 3 & conditioned: 12379-19240, Fig. 4a). This allowed us to rank sounds in terms of frequency range from more similar, to less similar to the conditioned sound. The most similar post-control sound shared a frequency band with the conditioned sound, but was discriminable by the frequency pattern. The sound cloud furthest away in frequency from the conditioned sound was used as the pre-control sound.

**Figure 4.**
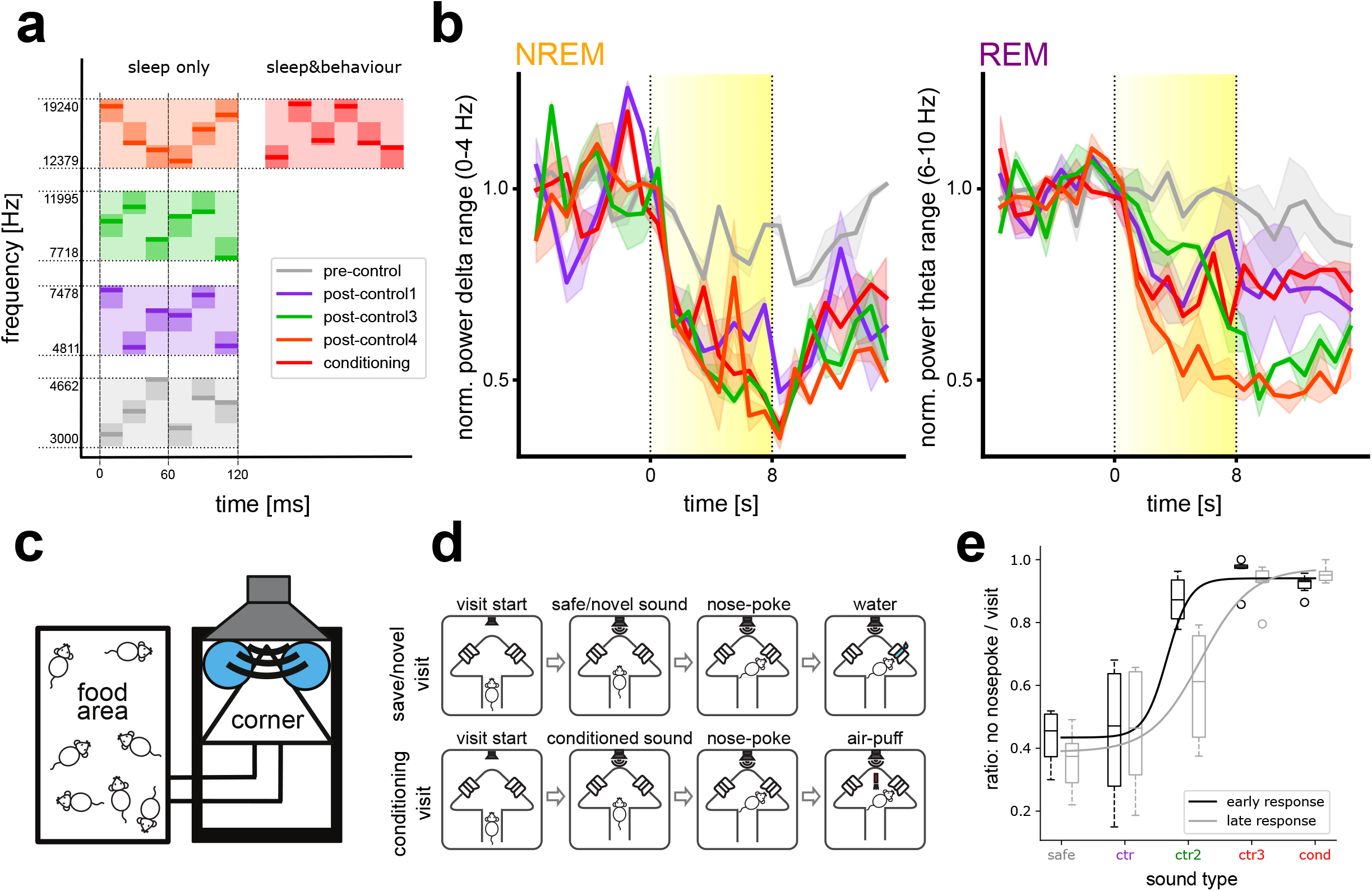
Sound generalization - EEG power changes to sound clouds. **(a)** Sound cloud architecture for all experimental sounds. Sound clouds were created pseudo randomly from three frequency blocks within the frequency range for each sound. The conditioning and post-control4 sound have the same frequency range, but differ in their frequency sequence. **(b)** Mean EEG power in delta (for NREM, left) and theta (for REM, right) ranges over time for learner animals (>80% correct trials on second day and > 15 trials in total). The five colours represent the five experimental sounds as in (a). Sound presentation is illustrated by a yellow block with sound intensity reflected in the colour intensity. (N = 2). **(c)** Audio box sketch with food area, tunnel, and water corner where the experimental sounds are presented. **(d)** Schematic of safe/novel and conditioned visits to corner. **(e)** Behavioural performance in Audiobox. Percentage avoidance of nose poking during a visit in the corner depending on sound played. The black line (first two days of exposure, early response) and the grey line (last two days of exposure late response) represent sigmoidal fits of the data. (N=6).

As found in the previous experiment, different sounds elicited different changes during NREM and REM sleep. In a 2-way ANOVA we found a main effect of state (F(1,458) = 28.52, p<0.001), sound (F(4,458) = 43.77, p<0.001) and time block (F(1,458) = 293.72 p<0.001). The pre-control sound had weak or no effect on ongoing oscillations and was significantly different from all other experimental sound during both NREM and REM sleep (Fig. 4b). Consistent with our previous results, we did not find a clear difference between the effects of the conditioned and post-control sound clouds during NREM sleep, and all sounds elicited strong effects that did not differ from each other (Fig. 4b). This time, however, we saw generalization also during REM sleep such that the post-control 3 sound cloud, in the same frequency band as the conditioned sound cloud, elicited the strongest changes and was significantly different from all other experimental sounds. Effects during REM sleep were generally weaker than in NREM sleep, however, consistent with the previous experiment.

To assess whether the strong generalization across sound clouds during sleep is due to an inability of the mice to discriminate between these sounds, we conducted a sound discrimination experiment in a different group of mice in the Audiobox, an automatic behavioural paradigm (Fig. 4c; explained in detail in de Hoz and colleagues^28^), using the same sounds. Mice were trained to access water in the water-corner through nose-poke holes (Fig. 4d). Whenever an animal entered the water-corner, a safe sound (the pre-control sound cloud) was played and the mouse was allowed to nose poke and thereby access water (Fig. 4d, safe visit). After this habituation phase, the conditioned sound cloud was introduced in 5% (then 10, then 20%) of water-corner visits. If the animal nose-poked during these visits, it would receive an air-puff and no access to water (Fig. 4d, conditioned visit). Mice typically avoided nose poking in only about a third of the visits, but quickly learned to avoid nose poking in most conditioned visits. When conditioned responses were stable, (>80% avoidance of nose pokes in conditioning visits and >50% nose poking in save visits) we sequentially introduced the other 3 novel sound clouds in 20% of visits each.

Our measure for deciding whether a novel sound was perceived more like the safe or conditioned sound was the ratio of nose pokes per visit. To account for possible learning effects, we separated the avoidance response into early and late response depending on the day of training, first two days and last two days respectively. Interestingly, the discrimination shifted subtly between early and late response and a 2-way ANOVA revealed a main effect of sound for the early (F(1,25) = 30.99, p<0.001) and late (F(1,25) = 23.44, p<0.001) responses. The post-control1, the closest in frequency to the safe sound, was consistently treated as safe by the animals (Fig. 4e). In. The post-control 2 sound, on the other hand, was categorized as unsafe during the early response (not significantly different from the conditioned sound, p = 0.9517), but later led to more nose pokes (significantly different from the conditioning sound, p<0.01). Overall, the data demonstrate that the sound clouds used are behaviourally discriminable and that the broad generalization across clouds observed in the sleep experiment is not the result of a lack of discriminability. Differences in sensory gating and generalization between sleep and wakening are not well understood and require further investigation.

## Discussion

We set out to investigate whether the meaning of a sound determines its influence on sleep-associated ongoing oscillations. We found that a frequency-modulated sound previously conditioned to an aversive outcome disrupted the EEG pattern during both NREM and REM sleep, leading to a decrease in state-dependent EEG power. These changes were observed at relatively low sound intensities of 30 to 40 dB SPL, not loud enough to elicit awakening or startle. That these changes are related to the aversive meaning of the sound is derived from the observation that a neutral FM sound presented in the same sleep bouts elicited weaker changes and only during NREM sleep. This latter effect was likely the result of generalization to the conditioning situation since a neutral sound presented before conditioning training began had no effect on either NREM or REM EEG power. We investigated the generalisation effect further using multiple neutral spectrotemporally rich sound clouds with varying frequency distance from the conditioned cloud. We found that generalization could be broad, such that sound clouds of frequencies that do not overlap with the conditioned sound also elicited changes in NREM EEG patterns. This was not related to the discriminability of the sound clouds, since mice trained to discriminate between these sounds could do so.

Thus, meaningful and neutral sounds affect distinct sleep stages differently. The effect that a sound has on sleep-associated ongoing oscillations does not depend only on its meaning but also on the sleep stage and day of conditioning. Thus, acquired behavioural relevance increases the potential of sounds to elicit changes during sleep in mice, which is in line with the findings in humans^29^, cats^30^and rats^31^. What do these EEG pattern changes mean? The EEG can give insight into global brain processes that are, nonetheless, dissociable from behavioural responses. For example, while we found clear changes to the EEG pattern upon sound presentation, we did not observe clear behavioural responses in the form of arousal. Early studies concluded that behaviourally significant sounds are more likely to arouse an animal than a neutral sound^30,31^. Yet none of these studies has provided a quantitative evaluation of EEG changes during sleep. Our study, by measuring changes in the EEG pattern using sound intensities that do not yet elicit measurable global arousal, provide a novel and more detailed perspective on auditory sensory gating during sleep. Nevertheless, the observed EEG pattern changes (decreased EEG delta power during NREM sleep and decreased theta band power during REM sleep) indicate a transition into lighter sleep or a more alert state. It is possible that the observed depression in EEG power resembles localised arousal that prepares the brain for transition into the waking state, while at the same time not yet disrupting sleep. Slow-wave activity (1-4 Hz EEG activity, SWA) has been suggested to be involved in sensory disconnection during sleep^32^, partially because associated off-states in neuronal firing may interfere with cortical signal propagation^33^. Thus, local or global depression of SWA could indicate a lowering of the waking threshold. For future studies, further physiological parameters like pupil size, whisker movements, heart rate or respiratory rate would offer useful measures in assessing the role of the observed EEG pattern changes in global arousal. Moreover, it might be interesting to further characterize NREM sleep in mice to identify similar sub-stages as are well known for humans^34^ or beginning to be recognised in rats^35^. That the effect of the conditioned sound on NREM EEG patterns is strong and reliable suggests, however, that the effect is not sensitive to subtle differences within NREM.

Another main finding was that the EEG pattern changed generally more during NREM compared to REM sleep. This is consistent with higher sensory thresholds during REM sleep^30,31^. We used different oscillation frequency ranges to measure the oscillatory power for NREM and REM, which makes the changes in power across both states not fully comparable. Yet, that the neutral sound presented during the conditioning training has an even weaker effect on REM sleep supports the idea that these two phases differ qualitatively in how they gate sensory information.

Besides the different potential to depress state dependent EEG power between conditioned and control sound we also found differences in the extinction of this effect over days. Even though the animals were conditioned every day before the recording session and the sound must have been highly familiar to the mouse, the decrease in power elicited by the conditioning sound remained profound and stable over the first 6 days during NREM sleep. In contrast, the control sound shows high variability during NREM sleep already during the first days of conditioning. This suggests that the generalization over frequency-modulated sounds was not influenced by the actual strength of the response to the conditioned sound itself. Interestingly, the effect elicited by the conditioned sound on REM sleep also weakened over days, again supporting our conclusion that the effect on different phases is qualitatively different.

Another interesting aspect of this experiment is the connection between wake behavioural performance and the potential of sounds to depress state-dependent EEG power. In animals that did not learn the task, sounds did not elicit EEG power changes as compared to ‘learning’ animals. This pronounced difference between learners and non-learners, observed already after the first conditioning session (before the behaviour could be classified as ‘non-learner’) allowed in fact predicting the behavioural performance of the animals on the next day.

Since we found that the effect of the conditioned sound generalized to the neutral sound during NREM sleep, we assessed the scope of this generalization now using spectrotemporally richer sounds in the form of sound clouds. We replicated the earlier finding of the control sound presented before the conditioning phase (pre-control sound) not eliciting EEG pattern changes. For the neutral sounds presented during the conditioning phase, the generalization was broad in that all sound clouds elicited some EEG power change during both NREM and REM, independently of their frequency. Moreover, while the sound cloud in a frequency range furthest from the conditioned sound elicited somewhat weaker effects, these were still clear during both NREM and REM. Interestingly, the neutral sound that was in the same frequency range as the conditioned sound, albeit with a different temporal architecture, elicited even stronger changes during REM sleep than the conditioned sound itself. The broader generalization was not caused by a lack of discriminability of the sounds, since another group of mice could distinguish between them in a sound discrimination task. These findings suggest first, that the generalization of the effect that sounds have on sleep patterns during REM might depend on the nature of the sounds themselves, as well as their familiarity. Since we had increased the number of neutral sounds presented during sleep, we did not have enough data to study the temporal development of the effect over days, when the effect of the neutral sounds on sleep might have weakened. Second, these findings suggest that the discriminability of sounds in the awake behaving animal does not automatically predict discriminability in the sleeping brain. Maybe sound features other than frequency, influence sensory gating in the sleeping brain (Chen et al., 2021). The criteria the sleeping brain applies in stimulus interpretation and the potential to shape generalisation during sleep remain to be addressed.

In conclusion, the meaning of a sound is a strong determinant of the effect this sound has on sleep-associated ongoing brain oscillations, even before any behavioural effect in the form of arousal can be detected. This effect is stronger, more persistent, and more generalizable to other sounds during NREM sleep. The paradigm presented here can be used as a model system to further explore the sensory gating mechanism and underlying circuits involved.

## Materials and Methods

### Animals

All experiments were aligned with the ethical regulations provided by the “Niedersachsisches Landesamt fur Verbraucherschutz und Lebensmittelsicherheit (LAVES)”. Project license number 33.9-42502-04-17/2571. The sound cloud experiment and the Audiobox discrimination-task experiment were conducted in Berlin, in accordance with the guidelines given by the “Landesamt fur Gesundheit und Soziales Berlin (LAGESO)” and were approved by this authority. Project license numbers G173/18, G140/19.

For awake EEG experiments combined with sound conditioning, we used eleven (9 for frequency-modulated experiment, 2 for sound cloud experiment) female C57BL/6J mice (*mus musculus*) from Janvier Labs. The animals were housed in standard plastic cages with a maximum of three littermates, *ad libitum* food and water access. Water restriction was introduced after one week of recovery following the surgery. Moreover, the animals were kept in a 12h/12h light/dark cycle (5:30 am/5:30 pm) in a temperature-controlled room (∼21°C). Cages were enriched with nesting material, a plastic hut and a paper roll. After surgery, the paper roll was removed and the plastic hut was turned on its head to remove possible obstacles from the implanted animals. Additionally, we used eight (6 included in analysis) female C57BL/6J mice for the discrimination task experiment in the Audiobox (TSE, Germany), where the mice lived for the duration of the experiment (see below). The amount of visits to the corner, nose pokes, consumed water and licks were analysed every day to ensure that all animals were drinking enough and in at least 50% of safe trials. Otherwise, mice were excluded from the experiment and kept under standard conditions (two out of eight animals).

### Surgery

For the implantation of the EEG drive and the EMG cable, the animal was anesthetized with either a reversible anaesthetic (**Göttingen:** Medetomidine [0.5 mg/Kg], Midazolam [5 mg/Kg], Fentanyl [0.05 mg/Kg]), or with a non-reversible one (**Berlin:** Ketamine [130 mg/Kg], Xylazine [10 mg/Kg]). Analgesia (**Göttingen:** Buprenorphine [0.1 mg/kg], **Berlin:** Carprofen [5 mg/Kg]) was injected subcutaneously as analgesic one hour before the end of the surgery. Analgesic was additionally provided subcutaneously every 24 hours for the following two days. Once anesthetized, the animal was fixed in a stereo tactic frame (Kopf Instruments) and the skin above the skull was cut prior subcutaneous injection of Lidocaine ([100 mg/Kg]). Holes were drilled (drill: Osada success 40; drill tip: Fine Science Tools, size 7) into the skull with the diameter of the silver painted screws for EEG and ground. Coordinates of EEG screw with respect to Bregma: anterior-posterior [-300], medial-lateral [-200]) for the screw positioned above the parietal cortex (Fig. 5a). To record the EMG the wire was sown through the neck muscle (*clavotrapezius*) and two knots were made to keep it in place. The screws and the EMG cable were connected to a custom-wired drive (see below). Once all connections were established, the drive was fixed to the skull with dental cement. **Göttingen:** anaesthesia was reversed using an antagonist: Flumazenil [0.5 mg/Kg], Atipamezole [2.5 mg/Kg]). The animals recovered for one week before the experiment begun.

**Figure 5.**
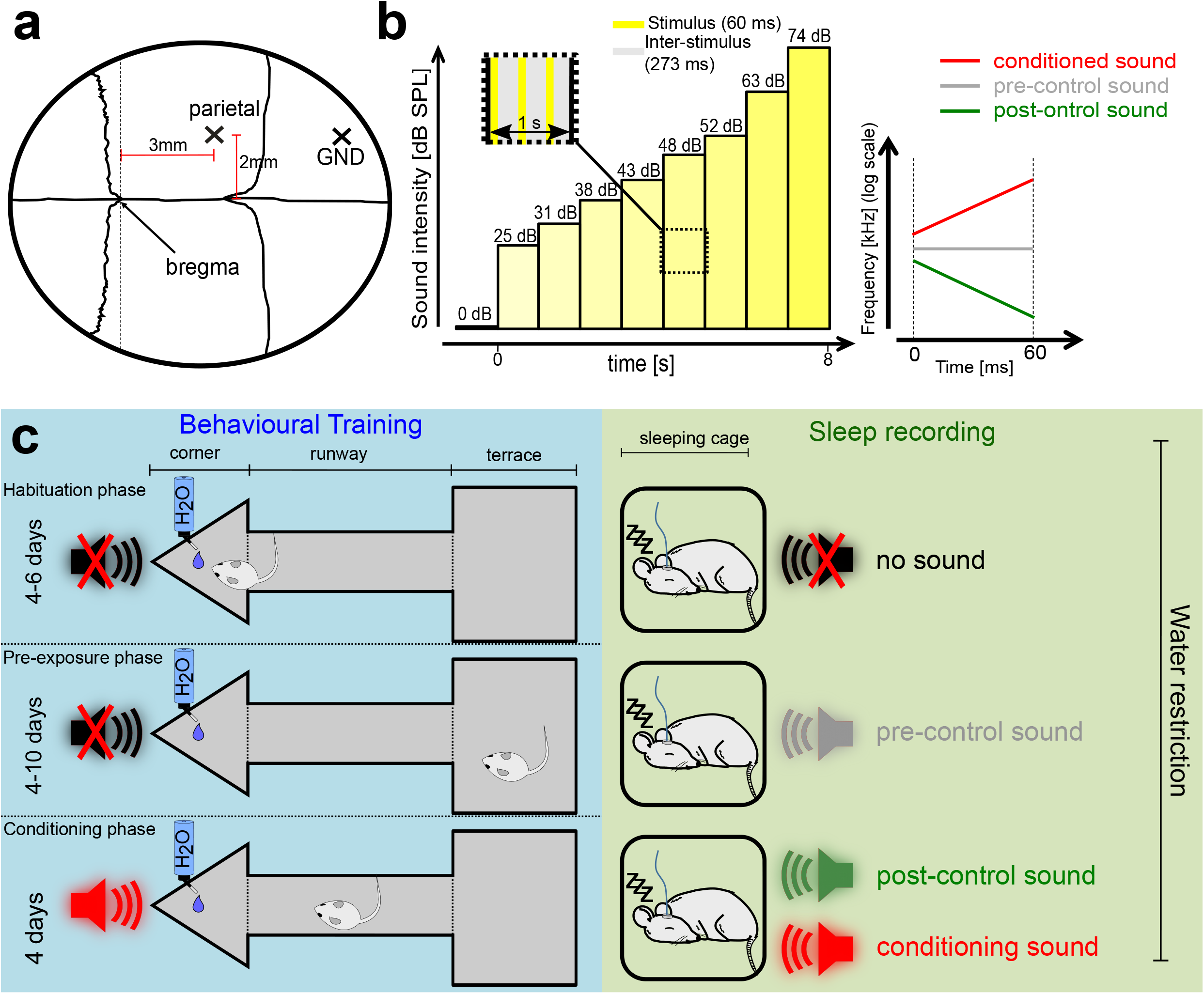
Experimental design. **(a)** Sketch of the mouse skull with implantation sites for parietal EEG and ground screw. **(b)** Left: Stimulus architecture over time. Insert shows stimulus and inter-stimulus time intervals within each intensity block. Eight intensity blocks (zero intensity excluded) each with three pulses adding up to 24 pulses over eight seconds. Right: Schematic of frequency modulated conditioned and post-control sound and pure tone pre-control sound. **(c)** Experimental Paradigm. Left (blue): Behavioural training on the audio terrace, consisting of corner, runway and terrace. Unlimited water access and no sounds in the corner during habituation and pre-exposure phase. During the conditioning phase, water access is granted for 20-40 seconds, before a conditioning trial starts. Right (green): Sound exposure during NREM and REM phases over the experimental stages.

The eight animals that were placed in the audiobox experiment underwent a minor surgery, where they were anesthetized with Ketamine-Xylazine (as above) and a small incision was made in the neck skin. A sterile transponder (PeddyMark, 12 mm × 2 mm or 8 mm × 1.4 mm ISO microchips, 0.1 gr in weight) was delivered under the skin and the skin closed with Histoacryl (B.Braun Surgical) to ensure a secure position of the transponder.

### In vivo recordings in awake animals

All awake electrical recordings were done with a chronically implanted micro drive (**Göttingen**: VersaDrive 4, Neurolynx, **Berlin**: 3D printed custom drive, Axona). EEG (stainless steel wire [203.2 μm coated diameter] with Teflon insulation, A-M Systems Inc.) and EMG (stainless steel wire [140 μm coated diameter] with PFA insulation, Science Products GmbH) wires were fed through this micro drive. The EEG was collected through a silver painted (Silberleitlack, Ferro GmbH) screw tightened in the skull over the parietal cortex (see Fig. 5a). During Recordings the micro drive was connected to an amplifier board (**Göttingen:** HS-36-Led, Neuralynx, USA; **Berlin:** RHD2132 16-Input Amplifier Board, Intan Technologies), which was connected to the acquisition system (**Göttingen:** Digital Lynx 4SX, Neuralynx, USA; **Berlin:** OpenEphys Acquisition board, OpenEphys). EEG and EMG signals were recorded (**Göttingen:** Cheetah Data Acquisition System software (Neuralynx, USA); **Berlin**: OpenEphys) at a sampling rate of 32 kHz or 30 kHz, respectively, and down sampled to 1000 Hz before analysis.

### Behavioural training on ‘audio-terrace’

The ‘audio-terrace’ is a metal construct on which the animal can move freely (see Fig. 5c, blue part). It consists of a ‘safe-terrace’, a connecting runway and a triangular corner where water access is given through two nose-poke holes. The sleeping cage of the animal was placed on the ‘safe-terrace’. In the sleeping cage, only an inverted hut and nest material was provided. The whole construct is elevated, placed on columns at a height of approximately 30 cm to restrict the animals from jumping off. In total four light barriers detected the movement of the animal over the terrace, whereby two were tracking the movement along the runway and one at each nose poke hole was tracking nose pokes. The training was automated via an analog-to-digital converter data acquisition system (NI SCB-68, National Instruments) controlled by custom written scripts (**Göttingen:** Matlab 2016b, Mathworks; **Berlin:** Python 3.6, Python Software Foundation, available at http://www.python.org). An ultrasonic speaker (Ultrasonic Dynamic Speaker Vifa, Avisoft) was placed above the nose poke corner during training on the ‘audio-terrace’ and over the sleeping cage during sleeping sessions (see Fig. 5c).

### Experimental protocol

Mice were 5 weeks of age upon arrival to the lab. After acclimation to the mouse room for at least one day, each mouse was handled 5 min daily to habituate the animal to the experimenter. After three days of this initial habituation, the implantation took place. After the surgery, the mice were allowed to recover for one week, in which they were still handled (five minutes daily). Then the experiment began. Within the first phase (habituation phase, compare Fig. 5c) the animal was trained for 20 min per day on the ‘audio-terrace’ and afterwards placed in the sleeping cage for a recording session of approximately 1.5 hours. On the audio-terrace, the animal was trained to access water by poking the nose-poke-holes in the corner. During the sleep sessions, the animals habituated to being connected to the recording system and undisturbed natural sleep was recorded. Starting with the pre-exposure phase, the training protocol on the terrace was not changed, but during sleep sessions, the pre-control sound was played to habituate the animals to being exposed to sounds while sleeping. Responses to the pre-control sound were recorded for two distinct sleep states (NREM & REM sleep). After successful training (min. 500ml water consumed during training), conditioning was introduced. During conditioning, the protocol on the audio-terrace included the timing of the start of a corner visit, and the conditioned sound was played randomly 20 to 40 s after the visit started. Each time the conditioned sound was played (i.e. the mouse spent enough time in the corner to trigger the sound) was counted as a trial. The conditioned sound was presented twice before an air puff and a loud sound were administered as aversive stimuli. If the mouse left the corner while the conditioned sound was still playing, the sound stopped and the trial was counted as successful, hence none of the aversive stimuli were administered. If the mouse did not escape on time, the conditioned sound was followed by punishment and the trial was counted as not successful. In Addition, the conditioned sound started to play again, continuing this loop until the mouse escaped. Mice rarely received more than two punishments per trial. Each time the mouse visited and left the corner, before the randomised timer triggered the beginning of the conditioned sound, was not considered a trial.

### Acoustic stimulation

For our experiment, we generated three to five (depending on experiment) sounds for every mouse. We used custom scripts (The Mathworks, Matlab, 2016b, or Python 3.6, Python Software Foundation, available at http://www.python.org). The sounds were created with a sampling rate of 200 kHz.

#### Logarithmically frequency modulated sounds (chirps)

The pre-control sound was a pure tone with a frequency in the centre of the other two experimental sounds and was selected from a range of frequencies (10-24 kHz). The conditioned sound was a frequency-modulated sound that started 500 Hz above the pre-control sound and went up at least two octaves at a rate of 50 octaves/second (Fig. 5b, right). The post-control sound was also frequency modulated, but the downward mirror image of the conditioned sound. This results in relatively similar initial frequency for the conditioned and post-control sounds and in increasing frequency differences over time.

#### Sound clouds

Sound cloud is a term to describe a type of auditory stimulus characterized by multiple tones within a defined frequency range. The tones are pure tones, but due to their randomized nature, sound clouds present auditory stimuli that have increased complexity compared to continuous frequency modulated sounds and pure tones. Due to their changing frequency, sound clouds are a type of frequency modulated sound, but differ from those used prior as they are a combination of separate pure tones rather than a continuous stimulus changing on a logarithmic scale. The sound architecture of the sound clouds used is visualized in Fig. 4a. Pre-control (3000 – 4662 Hz), post-control1 (4811 – 7478 Hz), post-control2 (7718 – 11995 Hz), post-control3 (12379 – 19240 Hz), and conditioned (12379 – 19240 Hz) were the five sounds used. Each sound cloud has an overall defined frequency range, or a ‘frequency window’. This window was further divided into three more narrow frequency ranges. Six pure tones were randomly selected within these three ranges, such that two tones were found within each range. These six tones were concatenated. As depicted, this overall frequency range is divided in half, and within each half, no tone is in the same frequency range as the one previously.

All sounds were played in a fixed scheme with intensities ranging from 24.6 to 73.5 dB in eight steps (Fig. 5b, left). Additionally, also a sequence with three repetitions of zero dB was played at the beginning of every sound to control for possible artefacts of the speaker. Sounds were played with a free-field ultrasonic speaker (Ultrasonic Dynamic Speaker Vifa, Avisoft) through an USB audio-interface (Octa-capture, Roland) and an amplifier (Portable Ultrasonic Power Amplifier, Avisoft). Sound triggers were sent simultaneously to the I/O board of the acquisition system. Sound intensity calibration was done with a calibrated microphone (Prepolarized Free-field 1/2’’ Microphone Type 4950) and a handheld analyser (Handgehaltener Analysator Typ 2250-L, Brüel & Kjær) measuring intensities for an eight kHz pure tone. All experiments were performed in a sound-attenuated and anechoic chamber.

### Behavioural training in Audiobox

The Audiobox is an automated system to conduct unsupervised learning experiments. For a more in depth explanation and sketch see ^28,36^. Briefly, the system consisted of two major parts, an experimental chamber and a home cage, which were connected via a tunnel. Food was provided *ad libitum* in the home-cage as well as nesting material, a hut and paper rolls, as environmental enrichment. To access water the mice had to follow a corridor to enter the drinking corner. This corner was located in a sound attenuated box and registered visits from individual animals by reading their unique transponders, which each animal was carrying underneath the neck skin. The corner had two nose poke holes, which could be closed or opened and when mice poked during safe visits the holes would open and water access was granted. The animals were housed in a group of eight animals and could perform ad libitum on their own schedule. The amount of visits to the corner, nose pokes and licks was analysed every day to ensure that all animals were drinking in at least 50% of safe trials, as was the case for all mice. During habituation all visits were safe, meaning that every time the mice entered the corner the save sound was played and poking the nose poke holes would result in water access. After all animals were familiar with the drinking system, the conditioned sound was introduced in 5% of visits. In conditioning visits, the conditioned sound was played and nose poking was punished by no water access and an air puff. During the following days the percentage of the conditioning visits was elevated to 20% or trials. After the stabilization of responses to the conditioned (nose poking during maximal 10% of conditioning trials) and save sound (nose poking during at least 50% of save trials), we started to introduce in total three novel sounds. In visits with novel sound, the corner was configured as for safe visits, meaning that when the mice nose poked they could access the water and no form of aversive stimulus was administered. We calculated the response of the animals to each of the experimental sounds by checking the percentage of visits in which there was a nose poke for each of the experimental sounds.

### Data analysis and statistics

The analysis and statistics of the electrophysiology and behavioural data was done in a Matlab/Python environment. For the sleep analysis, the power was extracted from the EEG-trace with the fast-Fourier transformation for frequencies of 0-4 Hz for NREM sleep and 6-10 Hz for REM sleep in a time window of 1 second for the 8 seconds before, during sound presentation and the 8 seconds afterwards. The resulting power data was normalized by the mean power across the 8 seconds prior to sound presentation. The statistical analysis, for all statistics reported, was done using multi-comparison ANOVA.

## Data availability

The datasets generated during and/or analysed during the current study are available from the corresponding author on reasonable request.

## Acknowledgements

We are grateful to Markus Krohn and his team at the MPI-EM workshop for technical support and the construction of the audio terrace.

## Author contributions

Conceptualization: L.M., L.d.H.

Data collection: P.v.K., L.M., Z.K.

Data curation: P.v.K., L.d.H.

Formal analysis: P.v.K., L.M., L.d.H.

Methodology: P.v.K., L.M., L.d.H.

Project administration: L.d.H.

Supervision: L.d.H.

Validation: P.v.K., L.d.H.

Visualization: P.v.K., L.d.H.

Writing – original draft: P.v.K., L.d.H.

Writing – review & editing: P.v.K., L.M., Z.K., L.d.H.

## Declaration of interests

The authors declare no competing interests.

## Notes

### Competing Interest Statement

The authors have declared no competing interest.

